# Satellite DNA-containing gigantic introns in a unique gene expression program during *Drosophila* spermatogenesis

**DOI:** 10.1101/493254

**Authors:** Jaclyn M Fingerhut, Jessica V. Moran, Yukiko M Yamashita

## Abstract

Intron gigantism, where genes contain megabase-sized introns, is observed across species, yet little is known about its purpose or regulation. Here we identify a unique gene expression program utilized for the proper expression of genes with intron gigantism. We find that two *Drosophila* genes with intron gigantism, *kl-3* and *kl-5*, are transcribed in a spatiotemporal manner over the course of spermatocyte differentiation, which spans ~90 hours. The introns of these genes contain megabases of simple satellite DNA repeats that comprise over 99% of the gene loci, and these satellite-DNA containing introns are transcribed. We identify two RNA-binding proteins that specifically localize to *kl-3* and *kl-5* transcripts and are needed for the successful transcription or processing of these genes. We propose that genes with intron gigantism require a unique gene expression program, which may serve as a platform to regulate gene expression during cellular differentiation.

## Introduction

Introns, non-coding elements of eukaryotic genes, often contain important regulatory sequences and allow for the production of diverse proteins from a single gene, adding critical regulatory layers to gene expression (Shaul, 2017). Curiously, some genes contain introns so large that more than 99% of the gene locus is non-coding. In humans, neuronal and muscle genes are enriched amongst those with the largest introns (Scherer, 2010). One of the best-studied large genes, Dystrophin, a causative gene for Duchenne Muscular Dystrophy, spans 2.2Mb, only 11kb of which is coding. A large portion of the remaining non-coding sequence is comprised of repetitive DNA-rich introns (Pozzoli et al., 2002). While intron size (‘gigantism’) is conserved between mouse and human, there is little sequence conservation within the introns, implying the functionality of intron gigantism (Pozzoli et al., 2003).

The *Drosophila* Y chromosome provides an excellent model for studying intron gigantism. Approximately 80% of the 40Mb Y chromosome is comprised of repetitive sequences, primarily satellite DNAs, which are short tandem repeats, such as (AATAT)_n_ (Figure 1A)(Carvalho, 2002; Hoskins et al., 2002; Lohe et al., 1993; Peacock et al., 1978). The *Drosophila* Y chromosome encodes fewer than 20 genes (Carvalho et al., 2015), six of which are collectively known as the ‘fertility factors’ (Brosseau, 1960; Gatti, 1983; Hazelrigg, 1982; Kennison, 1981). One of the fertility factors, *kl-3*, which encodes an axonemal dynein heavy chain (Carvalho et al., 2000; Goldstein et al., 1982; Hardy et al., 1981), spans at least 4.3Mb (Bonaccorsi et al., 1988; Gatti, 1983; Pimpinelli, 1986), while its coding sequence is only ~14kb (Figure 1A). This is due to the large satellite DNA rich-introns, some of which are megabases in size that comprise more than 99% of the *kl-3* locus. The other fertility factors (*kl-1, kl-2, kl-5, ks-1, ks-2*), have a similar gene structure, possessing large introns of repetitive satellite DNAs (Gatti, 1983). These six large Y chromosome genes are solely expressed during spermatogenesis (Bridges, 1916; Hardy et al., 1981; Marsh and Wieschaus, 1978).

**Figure 1.**
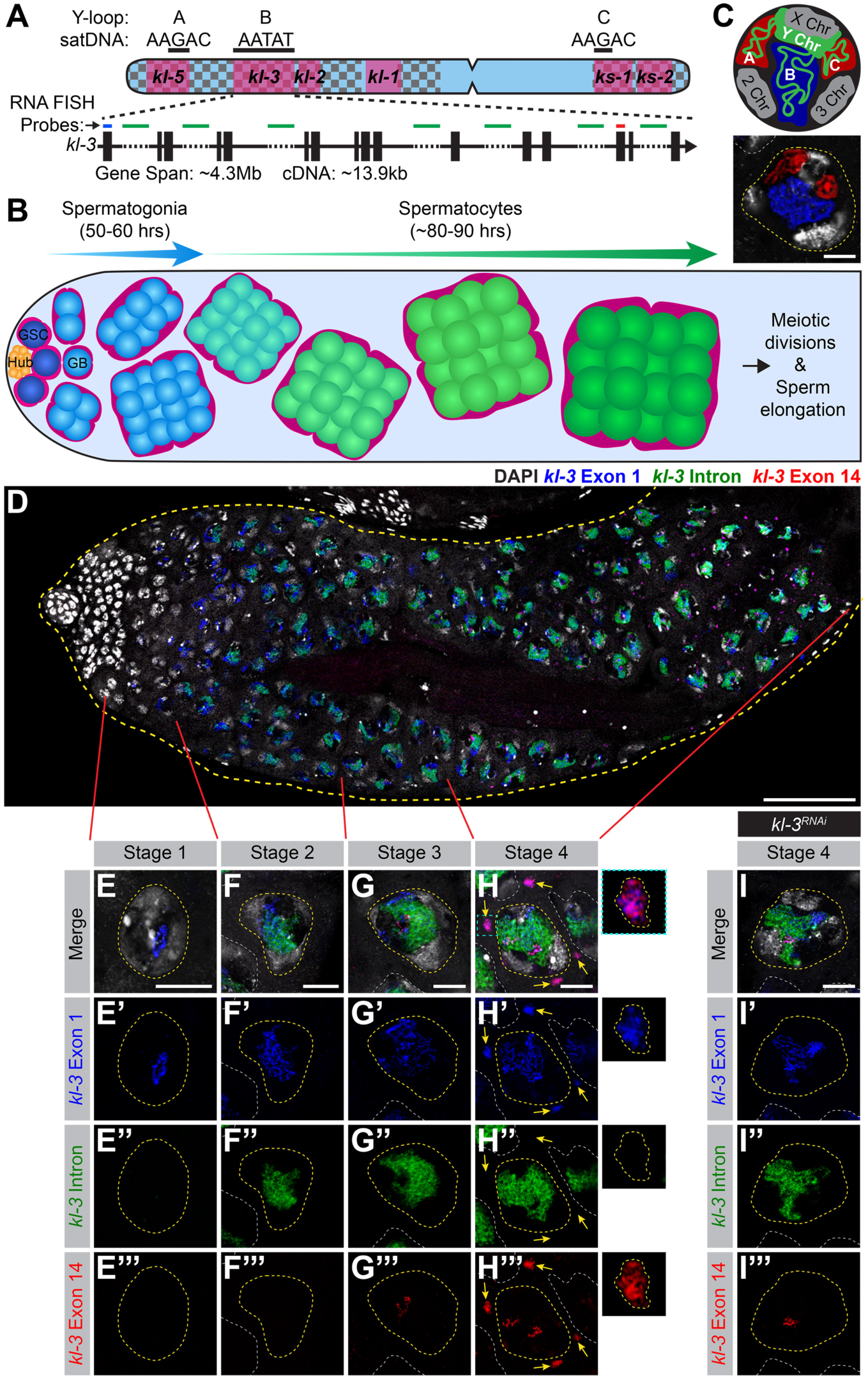
The Y-loop gene *kl-3* is expressed in a spatiotemporal manner during SC development. **(A)** Diagram of the *Drosophila* Y chromosome. Regions enriched for satellite DNA (checkered pattern), locations of the fertility factor genes (magenta) and the Y-loop forming regions (black bars) with associated satellite DNA sequences are indicated. Enlarged is a diagram of the Y-loop B gene *kl-3*. Exons (vertical rectangles), introns (black line), intronic satellite DNA repeats (dashed line) and regions of *kl-3* targeted by RNA FISH probes (colored bars). **(B)** Diagram of *Drosophila* spermatogenesis: GSCs (attached to the hub) produce mitotically-amplifying SGs, which become SCs. SCs develop over an 80–90 hour G2 phase before initiating the meiotic divisions. **(C)** Top: SC nucleus model showing the Y-loops in the nucleoplasm. DNA (white), Y chromosome (green), Y-loops A and C (red) and Y-loop B (blue). Bottom: RNA FISH for the Y-loop gene intronic transcripts in a SC nucleus. Y-loops are visualized using probes for Y-loops A and C (Cy3-(AAGAC)_6_, red) and Y-loop B (Cy5-(AATAT)_6_, blue). DAPI (white), SC nucleus (yellow dashed line) and nuclei of neighboring cells (white dashed line). Bar: 10μm. **(D-H)** RNA FISH to visualize *kl-3* expression in wildtype testes. Exon 1 (blue), *kl-3* intron (Alexa488-(AATAT)_6_, green), Exon 14 (red) and DAPI (white). **(D)** Apical third of the testis through the end of SC development (yellow dashed line). Bar: 75μm. **(E-H)** Single SC nuclei (yellow dashed line) at each stage of *kl-3* expression. Nuclei of neighboring cells (white dashed line) and cytoplasmic mRNA granules (yellow arrows). Bar: 10μm. Inset: *kl-3* mRNA granule (yellow dashed line). **(I)** RNA FISH against *kl-3* following *kl-3* RNAi (*bam-gal4>UAS-kl-3^TRiP.HMC03546^*). Single late SC nucleus (yellow dashed line) and nuclei of neighboring cells (white dashed line). Bar: 10μm.

In the *Drosophila* testis, germ cells undergoing differentiation are arranged in a spatiotemporal manner, where the germline stem cells (GSCs) reside at the very apical tip and differentiating cells are gradually displaced distally (Figure 1B) (Yamashita, 2018). GSC division gives rise to spermatogonia (SG), which undergo four mitotic divisions with incomplete cytokinesis to become a cyst of 16 SGs. 16-cell SG cysts enter meiotic S phase, at which point they become known as spermatocytes (SCs). SCs have an extended G2 phase, spanning 80–90 hours, prior to initiation of the meiotic divisions. During this G2 phase, the cells increase approximately 25 times in volume and the homologous chromosomes pair and segregate into individual chromosome territories (Figure 1C) (Fuller, 1993; McKee et al., 2012). During this period, SCs transcribe the majority of genes whose protein products will be needed for meiotic division and spermiogenesis (Gould-Somero and Holland, 1974; Olivieri and Olivieri, 1965; Schafer et al., 1995). Gene expression in SCs is thus tightly regulated to allow for timely expression of meiotic and spermiogenesis genes (White-Cooper and Caporilli, 2013).

It has long been known that three of the Y-chromosome-associated genes that contain gigantic introns (*kl-5, kl-3* and *ks-1*, Figure 1A) form lampbrush-like nucleoplasmic structures in SCs, named Y-loops [denoted as loops A (*kl-5*), B (*kl-3*), and C (ks-1), (Figure 1C, D)] (Bonaccorsi et al., 1988). Y-loop structures reflect the robust transcription of underlying genes, and have been observed across Drosophilids, including *D. simulans, D. yakuba, D. pseudoobscura, D. hydei* and *D. littoralis* (Hess, 1967; Piergentili, 2007). Much of the fundamental knowledge about Y-loops comes from *D. hydei*, which forms large, cytologically distinct Y-loops (Hackstein et al., 1982), leading to the discovery that these structures are formed by the transcription of large loci comprised of repetitive DNAs (Hochstenbach et al., 1994; Huijser et al., 1990; Trapitz et al., 1992; Vogt, 1986a; Vogt, 1986b; Wlaschek et al., 1988). Interestingly, in *D. pseudoobscura*, which contains a ‘neo-Y’ (not homologous to the ancestral Y chromosome), Y-loops are thought to be formed by Y-linked genes instead of by the *kl-3, kl-5* and *ks-1* homologs, which are on the autosomes (Chang and Larracuente, 2017), suggesting that Y-loop formation is a unique characteristic of Y-linked genes, instead of being a gene-specific phenomenon.

The transcription/processing of such gigantic genes/RNA transcripts, in which exons are separated by megabase-sized introns, must pose a significant challenge for cells. However, how genes with intron gigantism are expressed and whether intron gigantism plays any regulatory role in gene expression remain largely unknown. In this study, we began addressing these questions by using the Y-loop genes as a model, and describe the unusual nature of the gene expression program associated with intron gigantism. We find that transcription of Y-loop genes progresses in a strictly spatiotemporal manner, encompassing the entire ~90 hours of SC development: the initiation of transcription occurs in early SCs, followed by the robust transcription of the satellite DNA from the introns, with cytoplasmic mRNA becoming detected only in late SCs. We identify two RNA-binding proteins, Blanks and Hephaestus (Heph), which specifically localize to the Y-loops, and show that they are required for robust transcription and/or proper processing of the Y-loop gene transcripts. Mutation of the *blanks* or *heph* genes leads to sterility due to the loss of Y-loop gene products. Our study demonstrates that genes with intron gigantism require specialized RNA-binding proteins for proper expression. We propose that such unique processing may be utilized as an additional regulatory mechanism to control gene expression during differentiation.

## Results

### Transcription of a Y-loop gene, *kl-3*, is spatiotemporally organized

To start to investigate how the expression of Y-loop genes may be regulated, we sought to monitor their expression during SC development. In previous studies using *D. hydei*, when two differentially-labeled probes against two intronic repeats of the Y-loop gene *DhDhc7*(*y*) (homologous to *D. melanogaster kl-5*) were used for RNA fluorescent *in situ* hybridization (FISH), expression of the earlier repeat preceded that of the later repeat (Kurek et al., 1996; Reugels et al., 2000), leading to the idea that Y-loop genes might be transcribed as single, multi-megabase, transcripts. Consistently, Miller spreading of SC chromosome with transcripts still bound to DNA showed the long Y-loop transcripts (de Loos et al., 1984; Grond, 1983). However, transcription of exons was not visualized and extensive secondary structures were present in the Miller spreads, leaving it unclear whether the entire gene region is transcribed as a single transcript.

By using differentially-labeled probe sets designed for RNA FISH to visualize 1) the first exon, 2) the satellite DNA (AATAT)_n_ repeats found in multiple introns including the first (Bonaccorsi and Lohe, 1991; Lohe et al., 1993), and 3) exon 14 (of 16) of *kl-3* (Figure 1A, Supplementary file 1), we found that *kl-3* transcription is organized in a spatiotemporal manner: transcript from the first exon becomes detectable in early SCs, followed by the expression of the (AATAT)_n_ satellite from the introns, then finally by the transcript from exon 14 in more mature SCs (Figure 1D). These results suggest that transcription of *kl-3* takes the entirety of SC development, spanning ~90 hours. The pattern of transcription is consistent with the model proposed for Y-loop gene expression in *D. hydei:* the gene is likely transcribed as a single transcript that contains the exons and gigantic introns, although we cannot exclude the possibility of other mechanisms, such as the trans-splicing of multiple individually transcribed exons.

Based on the expression pattern of early exon, (AATAT)_n_ satellite-containing introns, and late exon, SC development can be subdivided into four distinct stages (Figure 1E-H). In stage 1, only exon 1 transcript is apparent (Figure 1E). In stage 2, the expression of intron transcript is detectable, while the signal from the first exon remains strong (Figure 1F). Stage 3 is defined by the addition of late exon signal, indicating that transcription is nearly complete (Figure 1G). Stage 4 is characterized by the presence of exon probe signals in granule-like structures in the cytoplasm (Figure 1H), which likely reflect *kl-3* mRNA localizing to ribonucleoprotein (RNP) granules, as they never contain intron probe signal. These granules are absent following RNAi-mediated knockdown of *kl-3* (*bam-gal4>UAS-kl-3^TRip.HMC03546^*, Figure 1I), confirming that they reflect *kl-3* mRNA. The same pattern of expression was seen for the Y-loop gene *kl-5* (see below, Figure 4B, C), suggesting that transcription of the other Y-loop genes proceeds in a similar manner.

Together, these results show that the gigantic Y-loop genes, including megabases of intronic satellite DNA repeats, are transcribed in a process that spans the entirety of SC development, culminating in the formation of mRNA granules in the cytoplasm near the end of the 80–90 hour meiotic G2 phase. This pattern of Y-loop gene transcription indicates that they are under precise regulation.

### Identification of genes that may regulate the transcription of the Y-loop genes

Considering the size of the Y-loop gene loci and their satellite DNA-rich introns, transcription of Y-loop genes likely utilizes unique regulatory mechanisms. To start to understand such a genetic program, we performed a screen (See Methods and Supplementary file 2). Briefly, a list of candidates was curated using a combination of gene ontology (GO) terms, expression analysis, predicted functionality and reagent availability, resulting in a final list of 67 candidate genes. Candidates were screened for several criteria including protein localization, fertility, and Y-loop gene expression. Among these, two genes, *blanks* and *hephaestus* (*heph*), exhibit localization patterns and phenotypes that reveal critical aspects of Y-loop gene regulation and were further studied. Several proteins, including Boule (Cheng et al., 1998), Hrb98DE (Lowe et al., 2014), Pasilla (Lowe et al., 2014; Redhouse et al., 2011) and Rb97D (Heatwole and Haynes, 1996), were previously shown to localize to the Y-loops but displayed no detectable phenotypes in Y-loop gene expression in SCs using RNAi-mediated knockdown and/or available mutants (Supplementary file 2), and were not further pursued in this study.

### Blanks and Heph are RNA binding proteins that specifically localize to the Y-loops and are required for fertility

Blanks, a RNA binding protein with multiple dsRNA binding domains, is primarily expressed in SCs in the testis. Blanks has been shown to be important for post-meiotic sperm development and male fertility (Gerbasi et al., 2011; Sanders and Smith, 2011). Blanks’, ability to bind RNA was found to be necessary for fertility (Sanders and Smith, 2011). In order to assess Blanks’ localization within the SC nucleus, testes expressing GFP-Blanks were processed for RNA FISH with probes against the intronic satellite DNA transcripts [(AATAT)_n_ for Y-loop B/*kl-3*, (AAGAC)_n_ for Y-loops *A/kl-5* and *C/ks-1* (Bonaccorsi et al., 1990)]. We found that GFP-Blanks exhibits strong localization to Y-loop B (Figure 2A).

**Figure 2.**
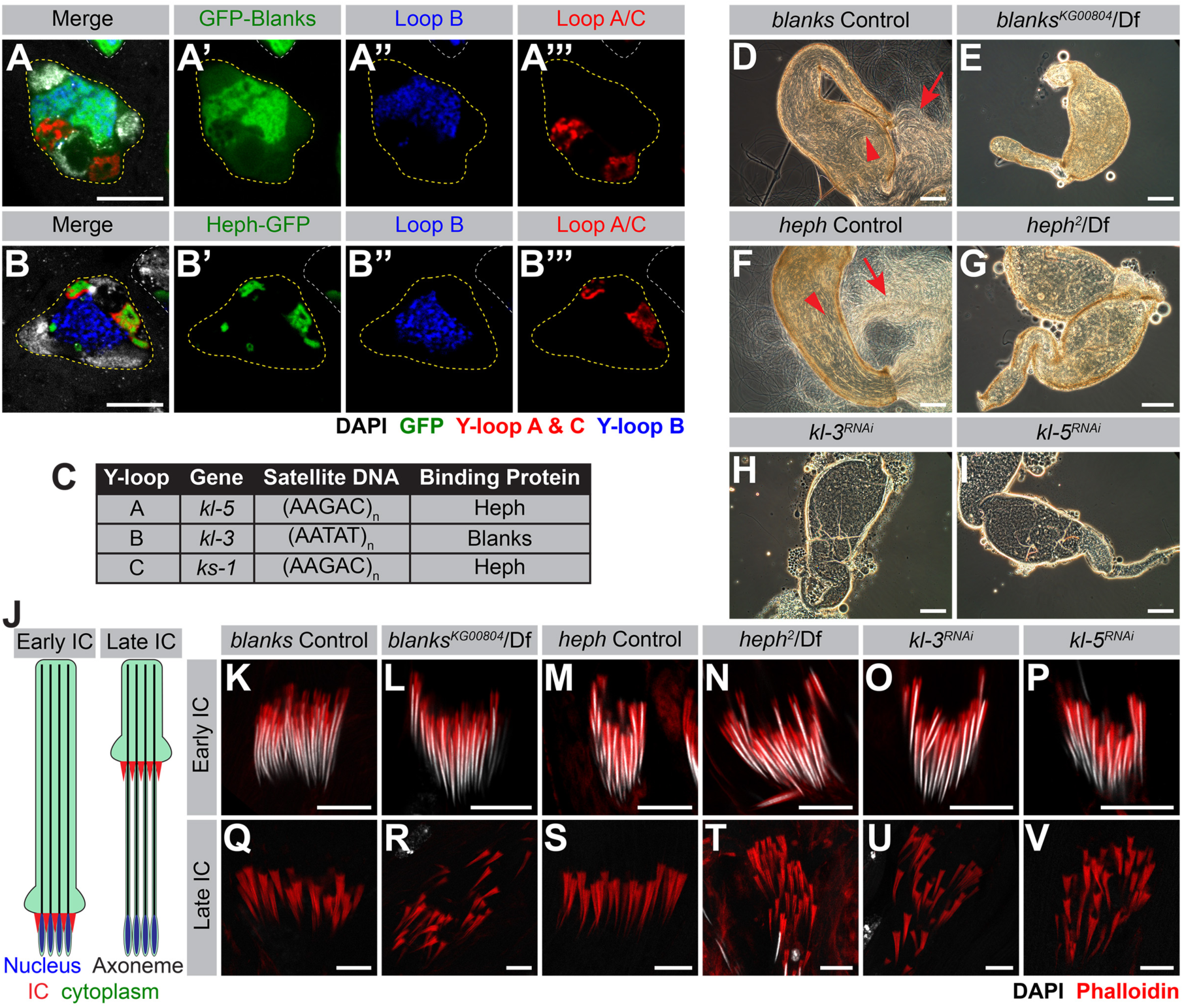
Blanks and Heph localize to the Y-loops and are required for fertility. **(A, B)** RNA FISH against the Y-loop gene intronic transcripts in flies expressing GFP-Blanks **(A)** or Heph-GFP **(B)**. Y-loops A and C (Cy3-(AAGAC)_6_, red), Y-loop B (Cy5-(AATAT)_6_, blue), GFP (green), SC nucleus (yellow dashed line) and nuclei of neighboring cells (white dashed line). Bar: 10μm. **(C)** Table listing Y-loop designation, associated gene, the satellite DNA repeats found within the introns and whether the Y-loop is bound by Heph or Blanks. **(D-I)** Phase contrast images of seminal vesicles in *blanks* controls **(C)**, *blanks^KG00084^*/Df **(D)**, *heph* controls **(E)**, *heph****^2^****/Df* **(F)**, *bam-gal4>UAS-kl-3^TRiP.HMC03546^* (**G**) and *bam-gal4>UAS-kl-5^TRip.HMC03747^* (**H**). Sperm within the seminal vesicle (red arrowhead) and extruded sperm (red arrow). Bar: 100μm. **(J)** Schematic of IC progression during individualization. Nucleus (blue), axoneme (black), ICs (red) and cytoplasm (green). **(K-V)** Phalloidin staining of early ICs **(K**-**P)** and late ICs **(Q-V)** in indicated genotypes. Phalloidin (actin, red) and DAPI (white). Bar : 10μm.

Heph, a heterogeneous nuclear ribonucleoprotein (hnRNP) homologous to mammalian polypyrimidine track binding protein (PTB), is a RNA-binding protein with multiple RNA recognition motifs (RRMs) that is expressed in the testis (Davis et al., 2002). Heph has also been implicated in post-meiotic sperm development and male fertility (Robida et al., 2010; Robida and Singh, 2003). By using a Heph-GFP protein trap (p(PTT-*GC*)*heph^CC00664^*) combined with RNA FISH to visualize the Y-loop gene intronic transcripts, we found that Heph-GFP localizes to Y-loops A and C (Figure 2B). It should be noted that the *heph* locus encodes 25 isoforms and the Heph-GFP protein trap likely represents only a subset of *heph* gene products. A summary of Y-loop designation, gene, intronic satellite DNA repeat, and binding protein is provided in Figure 2C.

We confirmed previous reports that *blanks* is required for male fertility (Gerbasi et al., 2011; Sanders and Smith, 2011). By examining the seminal vesicles for the presence of motile sperm, we found that seminal vesicles from control siblings contain abundant motile sperm (Figure 2D, 13% empty, 87% normal, n=46) while seminal vesicles from *blanks* mutants (*blanks^KG00084^*/Df(3L)BSC371) lack motile sperm (Figure 2E, 96% = empty, 4% greatly reduced, n=58). We also confirmed previous reports that *heph* is required for fertility (Castrillon et al., 1993; Robida et al., 2010). Seminal vesicles from *heph* mutants (*heph^2^*/Df(3R)BSC687) also lack motile sperm (Figure 2G, 100% empty, n=21), while those from control siblings contain motile sperm (Figure 2F, 5% empty, 95% normal, n=57).

Previous studies (Gerbasi et al., 2011; Robida et al., 2010; Sanders and Smith, 2011) reported that *blanks* and *heph* mutants are defective in sperm individualization, one of the final steps in sperm maturation, where 64 interconnected spermatids are separated by individualization complexes (ICs) that form around the sperm nuclei and migrate in unison along the sperm tails, removing excess cytoplasm and encompassing each cell with its own plasma membrane (Figure 2J) (Fabian and Brill, 2012). When the F-actin cones of IC were visualized by Phalloidin staining, it became clear that ICs form properly in all genotypes (Figure 2K-N), but become disorganized in the late ICs in *blanks* and *heph* mutants, preventing completion of individualization (Figure 2Q-T).

The sterility and individualization defects observed in *blanks* and *heph* mutants are reminiscent of the phenotypes observed in flies lacking axonemal dynein genes including *kl-5* and *kl-3*, the Y-loop A and B genes (Carvalho et al., 2000; Fatima, 2011; Gepner and Hays, 1993; Goldstein et al., 1982; Hardy et al., 1981; Timakov and Zhang, 2000). Upon RNAi mediated knockdown of *kl-3* and *kl-5* (*bam-gal4>UAS-kl-3^TRiP.HMC03546^* or *bam-gal4>UAS-kl-5^TRiR.HMC3747^*), motile sperm are not found in the seminal vesicles (Figure 2H, I, *kl-3:* 100% empty, n=81, *kl-5:* 94% empty, 6% reduced, n=50) and a scattering of late ICs is observed (Figure 2O-P, U-V). Based on these observations, we hypothesized that the sterility and ICs defects observed in *blanks* and *heph* mutants may arise due to failure in the expression of the Y-loop genes.

### blanks is required for transcription of the Y-loop B gene kl-3

As *blanks* was found to localize to Y-loop B, we first determined whether there were any overt defects in Y-loop B formation or *kl-3* expression in *blanks* mutants. To this end, we performed RNA FISH to visualize the Y-loop gene intronic transcripts in *blanks* mutants. Compared to control testes where intronic satellite DNA transcripts from all Y-loops become detectable fairly early in SC development and quickly reach full intensity (Figure 3A), the signal from the Y-loop B intronic transcripts remains faint in *blanks* mutants (Figure 3B). The expression of Y-loops A and C is comparable between control and *blanks* mutant testes (Figure 3A, B).

**Figure 3.**
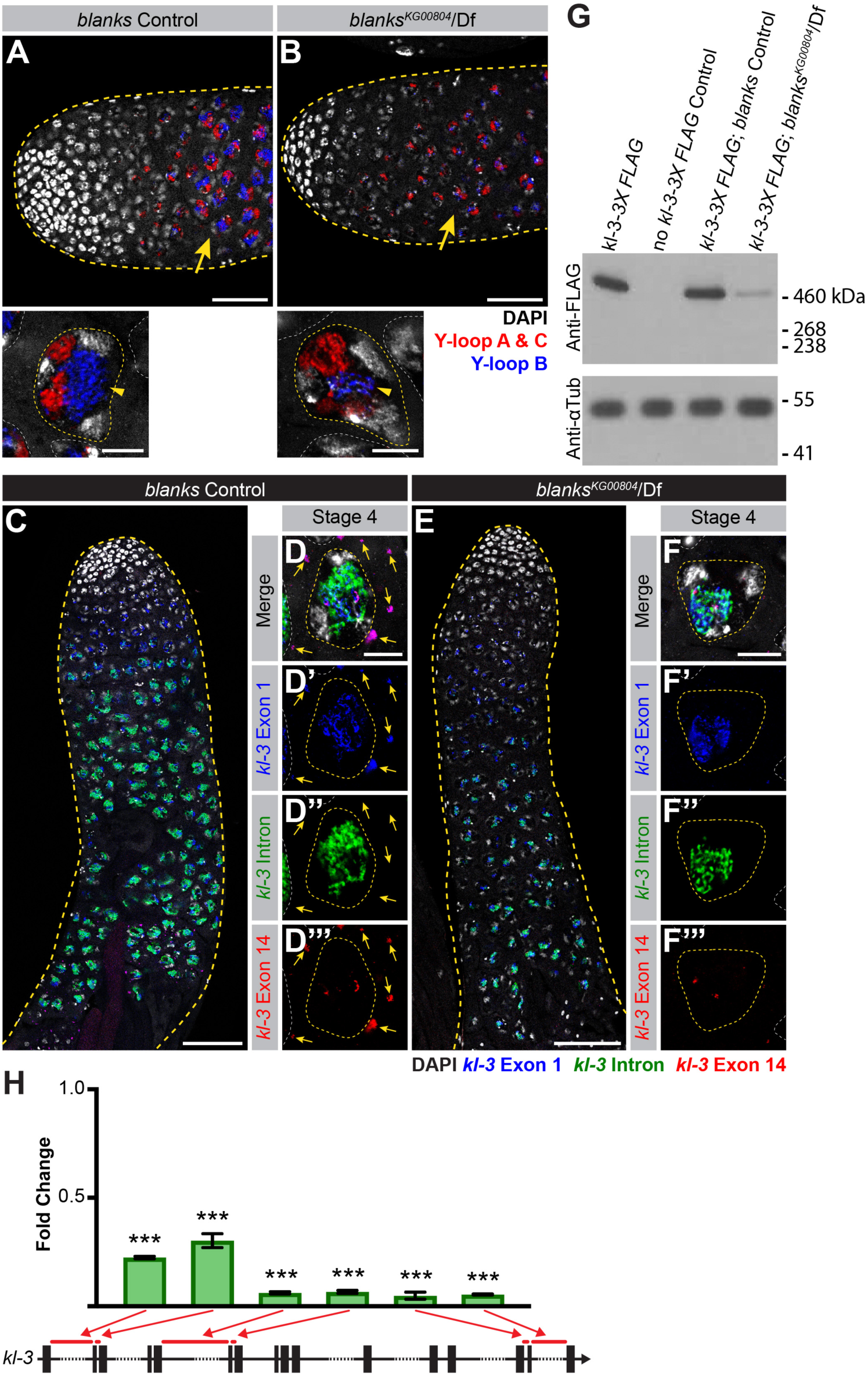
*blanks* is required for *kl-3* expression. **(A, B)** RNA FISH against the Y-loop gene intronic transcripts in *blanks* control **(A)** and *blanks^KG000804^*/Df **(B)**. Testis outline (yellow dashed line), Y-loops A and C (Cy3-(AAGAC)_6_, red), Y-loop B (Cy5-(AATAT)_6_, blue) and DAPI (white). Comparable stage SCs are indicated by yellow arrows. Bar: 50μm. High magnification images of single SCs at a comparable stage are provided below. SC Nuclei (yellow dashed line) and nuclei of neighboring cells (white dashed line). Bar: 10μm. **(C-F)** RNA FISH against *kl-3* in *blanks* control **(C, D)** and *blanks^KG00084^*/Df **(E, F)**. Exon 1 (blue), *kl-3* intron (Alexa488-(AATAT)_6_, green), Exon 14 (red) and DAPI (white). **(C, E)** Apical third of the testis through the end of SC development (yellow dashed line). Bar: 75μm. **(D, F)** Single late SC nucleus (yellow dashed line). Nuclei of neighboring cells (white dashed line) and mRNA granules (yellow arrows). Bar: 10μm. **(G)** Western blot for Kl-3–3X FLAG in the indicated genotypes. **(H)** RT-qPCR in *blanks^KG00084^/Df* for *kl-3* using the indicated primer sets. Primer locations are designated by red bars on the gene diagram. Data was normalized to GAPDH and sibling controls. Mean ±SD (p-value ***≤0.001 t-test between mutant and control siblings, exact p-values listed in Source Data 1)

In addition to a reduction in the expression of the intronic satellite DNA repeats of Y-loop B/*kl-*3, expression of *kl-3* exons is also reduced in *blanks* mutants. By performing RNA FISH using exonic and intronic (AATAT)_n_ probes for Y-loop B/*kl-3*, we found that *blanks* mutant flies display an overall reduction in signal intensity for both intronic satellite repeats and exons compared to controls (Figure 3C, E). Moreover, cytoplasmic *kl-3* mRNA granules are rarely detected in *blanks* mutants (Figure 3F). The same results are obtained following RNAi mediated knockdown of *blanks* (*bam-gal4>UAS-blanks^TRiP.HMS00078^*, Supplementary Figure 1). These results suggest that *blanks* is required for robust and proper expression of Y-loop B/*kl-3* and for the production of *kl-3* mRNA granules, likely at the transcriptional level. Consistently, we found that the amount of Kl-3 protein is greatly diminished in *blanks* mutants, confirming that *blanks* is required for proper expression of Y-loop B/*kl-3* (Figure 3G).

To obtain a more quantitative measure of *kl-3* expression levels in control and *blanks* mutant testes, we performed RT-qPCR. Primers were designed to amplify early (close to the 5’ end), middle, and late (close to the 3’ end) regions of *kl-3*. For each region, two sets of primers were designed: one primer set spanned a satellite-DNA-containing large intron and another spanned a normal size intron (Figure 3H, bars denote spanned intron, and Supplementary file 3). All primer sets show a detectable drop in *kl-3* mRNA levels in *blanks* mutants (Figure 3H). We noted a detectable drop between the early primer sets (~75% reduction in expression levels compared to controls) and the middle/late primer sets (~95% reduction in expression levels compared to controls), raising the possibility that *blanks* mutants may have difficulty transcribing this Y-loop gene soon after encountering the first satellite DNA containing gigantic intron. In summary, the RNA binding protein Blanks localizes to Y-loop B and allows for the robust transcription of the Y-loop B gene *kl-3*.

In contrast to Y-loop B/*kl-*3 expression, Y-loop *A/kl-5* expression appeared normal in *blanks* mutants. We designed RNA FISH probes against *kl-5* in the same manner as for *kl-3* (i.e. early exon, intron and late exon) (Figure 4A, Supplementary file 1). We found that transcription of Y-loop *A/kl-5* follows a spatiotemporal pattern similar to that of Y-loop B/*kl-3* (Figure 4B, C): early exon transcripts becomes detectable in early SCs while *kl-5* mRNA granules are not detected in the cytoplasm until near the end of SC development (Figure 4B, C). No overt differences are observed in *kl-5* expression in *blanks* mutants and *kl-5* mRNA granules are observed in the cytoplasm (Figure 4D, E). By RT-qPCR with primers for *kl-5* designed similarly as described above for *kl-3* (Figure 3H), we found a mild reduction in *kl-5* expression in *blanks* mutants (Figure 4F). However, considering the fact that the *kl-5* mRNA granule is correctly formed in *blanks* mutant testes (Figure 4E), this reduction may not be biologically significant. The mild reduction in *kl-5* transcript in *blanks* mutants could be an indirect effect caused by defective Y-loop B expression. Alternatively, it is possible that a small amount of (AATAT)_n_ satellite, which is predicted to be present in the last intron of *kl-5* (Bonaccorsi and Lohe, 1991; Hoskins et al., 2015), might cause this mild reduction in *kl-5* expression in *blanks* mutant testes.

**Figure 4.**
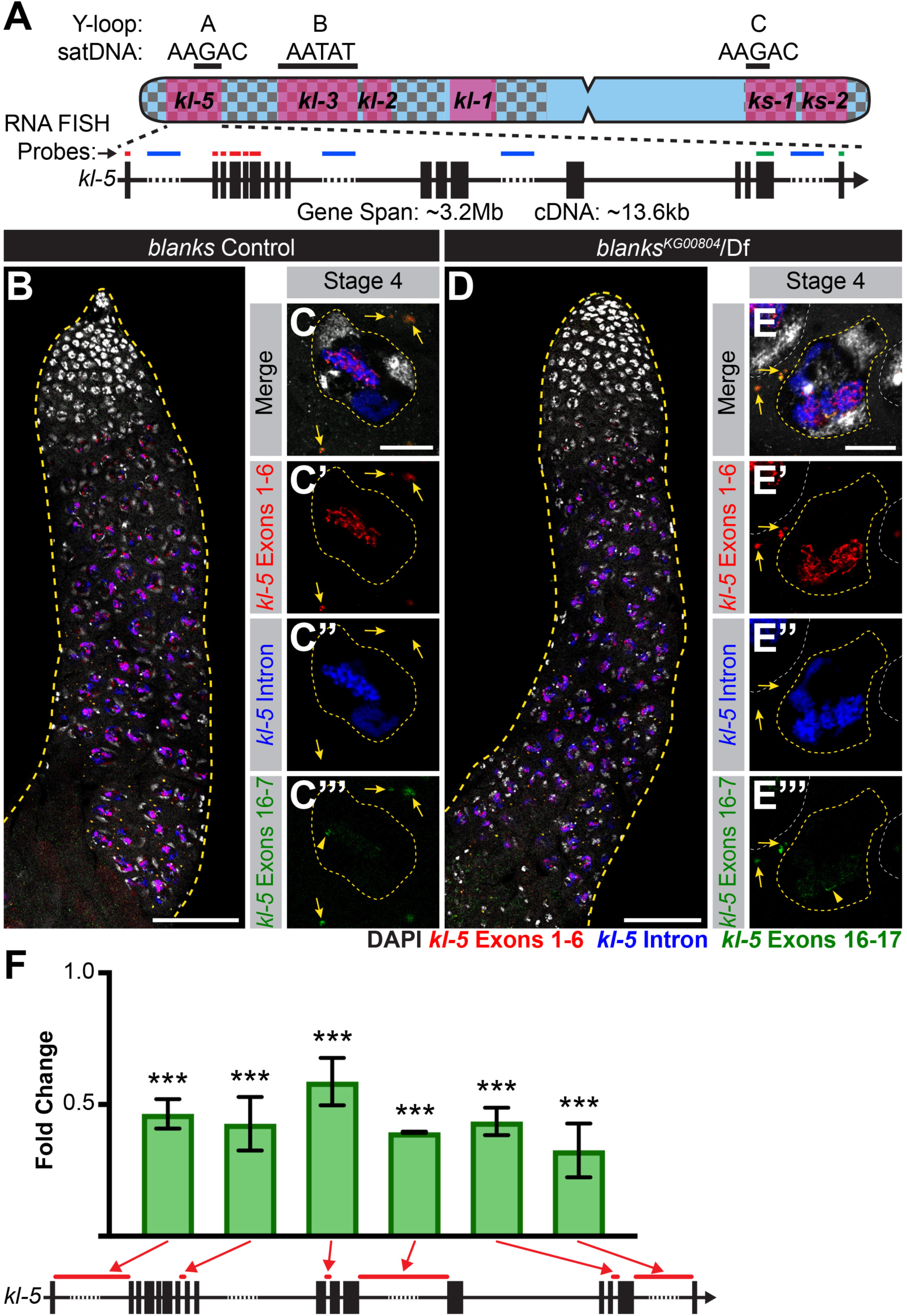
*blanks* is not required for *kl-5* expression. **(A)** Diagram of the Y-loop A gene *kl-5*. Exons (vertical rectangles), introns (black line) and intronic satellite DNA repeats (dashed line). Regions of *kl-5* targeted by RNA FISH probes are indicated by the colored bars. **(C-F)** RNA FISH against *kl-5* in *blanks* controls **(C, D)** and *blanks^KG00084^*/Df **(E, F)**. Exons 1–6 (red), *kl-5* intron (Cy5-(AAGAC)_6_, blue), Exons 16–17 (green, arrowhead indicates nuclear signal), DAPI (white). **(C, E)** Apical third of the testis through the end of SC development (yellow dashed line). Bar: 75μm. **(D, F)** Single late SC nucleus (yellow dashed line). Nuclei of neighboring cells (white dashed line) and mRNA granules (yellow arrows). Bar: 10μm. **(G)** RT-qPCR in *blanks^KG00084^*/Df for *kl-5* using the indicated primer sets. Primer locations are designated by red bars on the gene diagram. Data was normalized to GAPDH and sibling controls. Mean ±SD (p-value ***≤0.001, t-test between mutant and control siblings, exact p-values listed in Source Data 2).

### Blanks is unlikely to be a part of the general meiotic transcription program

It is well known that SCs utilize a specialized transcription program in order to transcribe the vast majority of genes required for meiosis and spermiogenesis (Lin et al., 1996; White-Cooper and Caporilli, 2013; White-Cooper et al., 1998). This program is executed by two groups of transcription factors: tMAC and the tTAFs. The tMAC (testis-specific meiotic arrest complex) complex has both activating and repressing activities and has been shown to physically interact with the core transcription initiation machinery (Ayyar, 2003; Beall et al., 2007; Doggett et al., 2011; Jiang, 2003; Jiang et al., 2007; Lu and Fuller, 2015; Perezgasga et al., 2004). The tTAFs (testis-specific TATA binding protein associated factors) are homologs of core transcription initiation factors (Chen et al., 2005; Hiller et al., 2004; Hiller et al., 2001; Laktionov et al., 2014). tMAC and the tTAFs function cooperatively to regulate meiotic gene expression. To examine whether *blanks* is part of this established meiotic transcription program, we examined the expression of *fzo* and *Dic61B*, known targets of the SC-specific transcriptional program (Laktionov et al., 2014; White-Cooper et al., 1998), which are located on autosomes and do not have gigantic introns (Supplementary file 1). In contrast to mutants for the tMAC component *aly* (*aly^2/5P^*), which has drastically reduced levels of *fzo* and *Dic61B* transcripts, the expression of these genes is not visibly affected in *blanks* mutants (Supplementary Figure 2), suggesting that *blanks* is not a part of the SC-specific transcriptional program involving tTAF and tMAC. Instead, *blanks* is likely uniquely involved in the expression of the Y-loop genes.

### Heph is required for processing transcripts of the Y-loop A gene *kl-5*

As Heph-GFP localized to Y-loops A and C, we first examined whether Y-loops A and C displayed any overt expression defects in *heph* mutants as seen in *blanks* mutants (Figure 3B). When we performed RNA FISH to visualize the Y-loop gene intronic transcripts in *heph* mutants, the overall expression levels of both (AAGAC)_n_ and (AATAT)_n_ satellites appear unchanged between control and *heph* mutant testes (Figure 5A, B). However, we noted that the morphology of Y-loops A and C is altered in *heph* mutants, adopting a less organized, diffuse appearance (Figure 5B), whereas all Y-loops in control SCs show characteristic thread-like or globular morphologies (Figure 5A). Y-loop B appears unchanged between controls and *heph* mutants (Figure 5A, B). These results indicate that *heph* may be important for structurally organizing Y-loop A and C transcripts, without affecting overall transcript levels.

**Figure 5.**
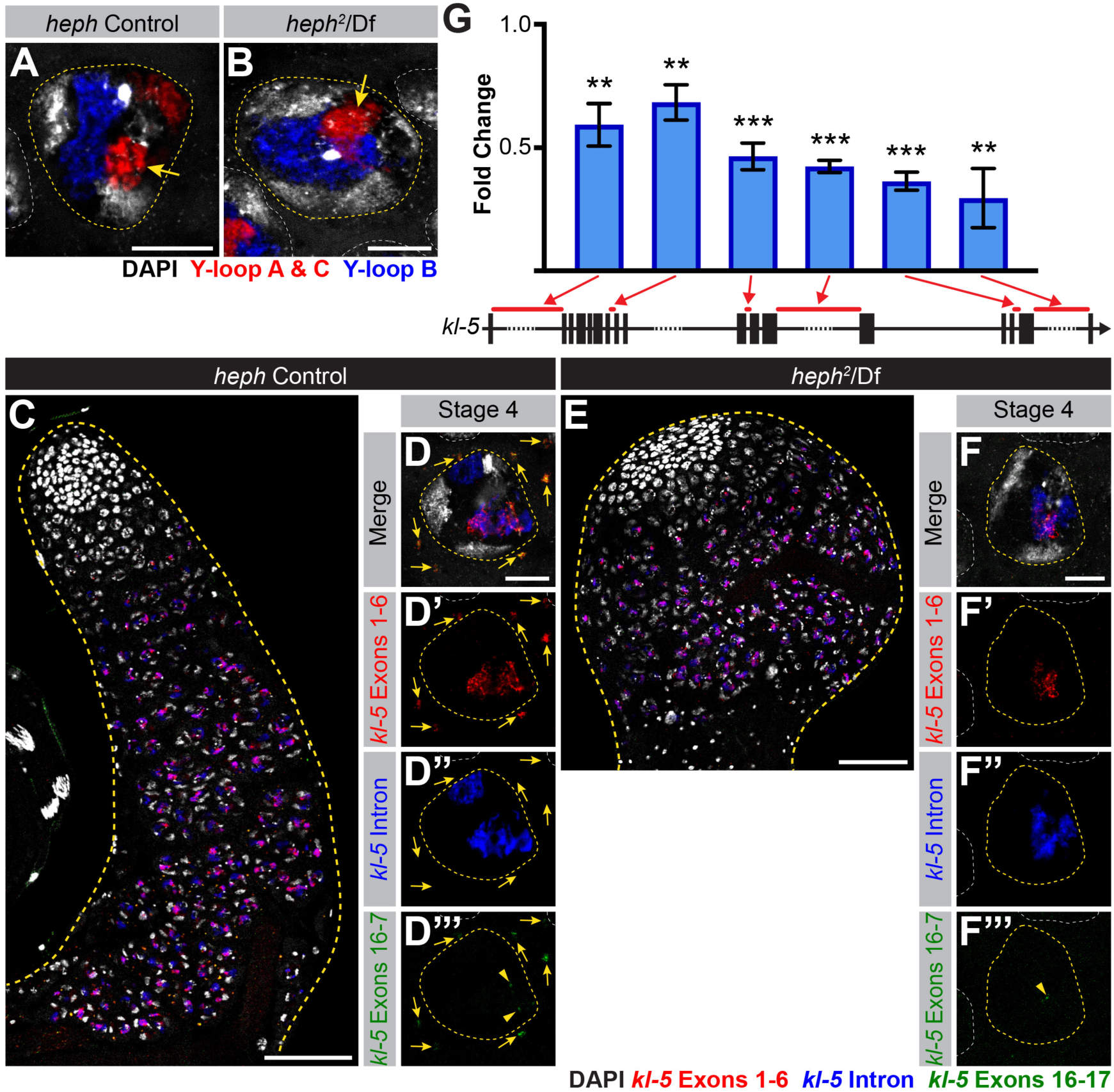
*heph* is required for *kl-5* mRNA production. **(A, B)** RNA FISH against the Y-loop gene intronic transcripts in *heph* controls **(A)** and *heph^2^*/Df **(B)**. Single late SC nuclei (yellow dashed line), nuclei of neighboring cells (white dashed line), Y-loops A and C (Cy3-(AAGAC)_6_, red), Y-loop B (Cy5-(AATAT)_6_, blue) and DAPI (white). Bar: 10μm. **(C-F)** RNA FISH against *kl-5* in *heph* controls **(C, D)** and *heph^2^*/Df **(E, F)**. Exons l-6 (red), *kl-5* intron (Cy5-(AAGAC)_6_, blue), Exons 16–17 (green, arrowhead indicates nuclear signal) and DAPI (white). **(C, E)** Apical third of the testis through the end of SC development (yellow dashed line). The bulbous shape of *heph^2^*/Df is a known phenotype that can occur with this allele (Castrillon et al., l993). Bar: 75μm. **(D, F)** Single late SC nuclei (yellow dashed line). Nuclei of neighboring cells (white dashed line) and mRNA granules (yellow arrows). Bar: 10μm. **(G)** RT-qPCR in *heph^2^*/Df for *kl-5* using the indicated primer sets. Primer locations are designated by red bars on the gene diagrams. Data was normalized to GAPDH and sibling controls. Mean ±SD (p-value **≤0.01, ***≤0.001, t-test between mutant and control siblings, exact p-values listed in Source Data 3).

We next examined expression pattern of *kl-5* exons together with the Y-loop A/C intronic satellite [(AAGAC)_n_] as described in Figure 4. Overall expression levels of *kl-5* appear to be unaltered in *heph* mutant testes (Figure 5C, E). However, in contrast to control testes (Figure 5D), *heph* mutant testes rarely have cytoplasmic *kl-5* mRNA granules in late SCs (Figure 5E), suggesting that *heph* mutants affect *kl-5* mRNA production without affecting transcription in the nucleus. *heph* mutants may be defective in processing the long repetitive regions of transcripts to generate mRNA (e.g. splicing, mRNA export). Similar to *blanks* mutants, *heph* mutants do not affect the expression of *fzo* or *Dic61B* (Supplementary Figure 2), indicating that *heph* is not a member of the more general meiotic transcription program. Instead, *heph*, like *blanks*, appears to specifically affect the expression of Y-loop *A/kl-5* to which it localizes.

RT-qPCR showed that *heph* mutants only exhibit a moderate reduction in *kl-5* expression (Figure 5G), which is in accordance with the RNA FISH results described above. A similar moderate reduction in *kl-5* mRNA is observed in *blanks* mutants (Figure 4F), which do not affect *kl-5* mRNA granule formation. Thus, it is unlikely that the reduction in *kl-5* expression levels alone causes the lack of *kl-5* mRNA granules in *heph* mutant SCs. Instead, we postulate that mRNA granule formation is dependent on proper processing of primary transcripts, which may be defective in *heph* mutants.

Surprisingly, we found that *kl-3* mRNA granules are also absent in *heph* mutants, although Y-loop B/*kl-3* expression levels in the nucleus appear to be unaffected (Figure 6A-D). RT-qPCR showed a similar moderate reduction in *kl-3* mRNA in *heph* mutants (Figure 6E) as was observed in *kl-5* mRNA (Figure 5G). Consistent with the absence of cytoplasmic *kl-3* mRNA granules, Kl-3 protein levels are dramatically reduced in *heph* mutant testes (Figure 6F). This is unexpected as Heph protein does not localize to Y-loop B (Figure 2B) or affect Y-loop B morphology (Figure 5A, B). It is possible that some of the predicted 25 isoforms of Heph are not visualized by Heph-GFP, and these un-visualized isoforms might localize to and regulate Y-loop B/*kl-*3 expression. Alternatively, this may be an indirect effect of defective Y-loop A and C expression and/or structure.

**Figure 6.**
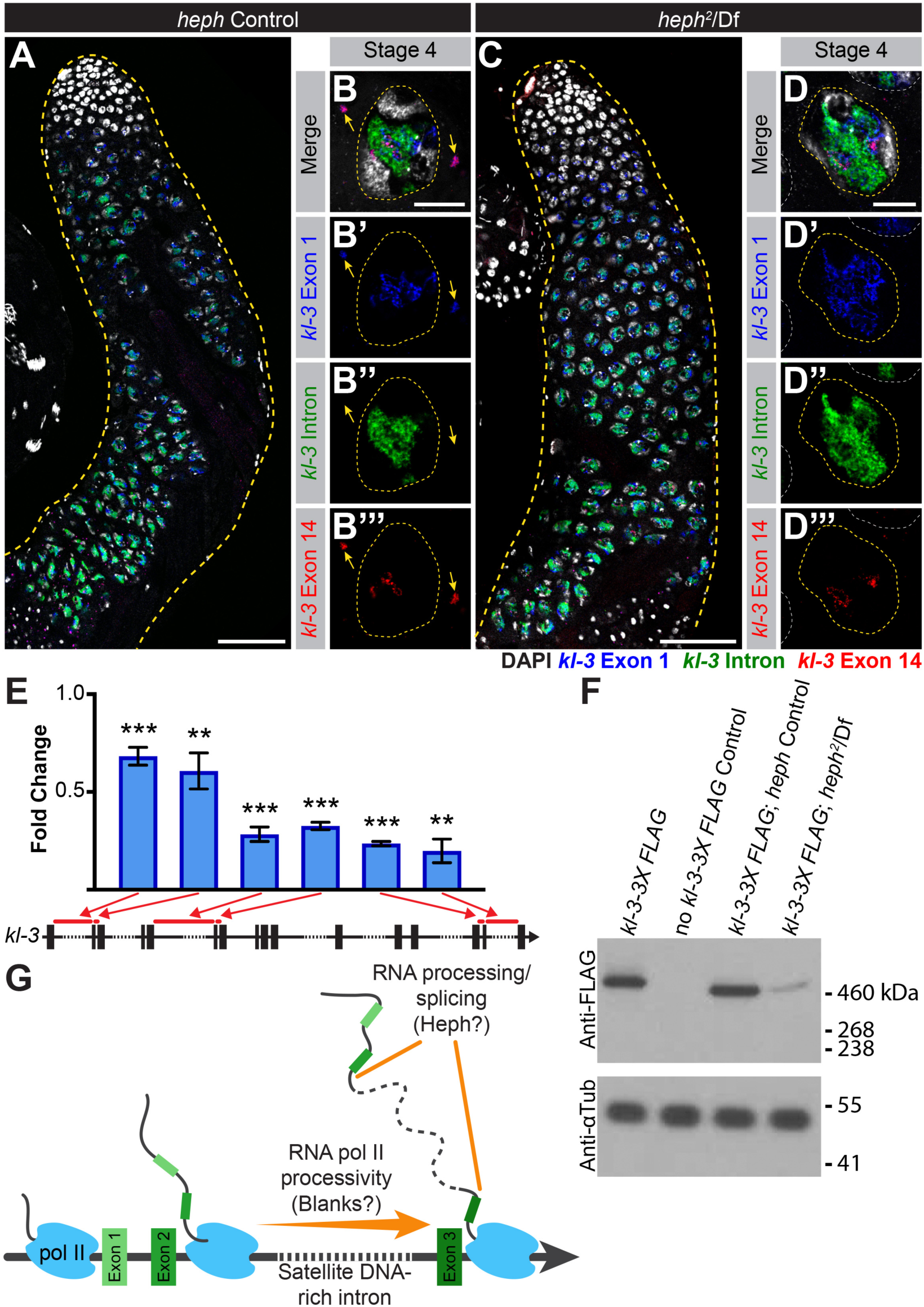
*kl-3* expression is affected in *heph* mutants. **(A - D)** RNA FISH against *kl-3* in *heph* controls **(A, B)** and *heph^2^*/Df **(C, D)**. Exon 1 (blue), *kl-3* intron (Alexa488-(AATAT)_6_, green), Exon 14 (red) and DAPI (white). **(A, C)** Apical third of the testis through the end of SC development (yellow dashed line). Bar: 75μm. **(B, D)** Single late SC nuclei (yellow dashed line). Nuclei of neighboring cells (white dashed line) and mRNA granules (yellow arrows), Bar: 10μm. **(E)** RT-qPCR in *heph^2^*/Df for *kl-3* using the indicated primer sets. Primer locations are designated by red bars on the gene diagrams. Data was normalized to GAPDH and sibling controls. Mean ±SD (p-value **≤0.01 ***≤0.001, t-test between mutant and control siblings, exact p-values listed in Source Data 4). **(F)** Western blot for Kl-3-3X FLAG in the indicated genotypes. **(G)** Model for the Y-loop gene transcriptional program.

Taken together, our results show that Blanks and Heph, two RNA binding proteins, are essential for the expression of Y-loop genes, but are not members of the more general meiotic transcription program. As Y-loop genes are essential for sperm motility and fertility, the sterility observed in *blanks* and *heph* mutants likely stems from defects in Y-loop gene expression. Blanks and Heph highlight two distinct steps (transcriptional processivity and RNA processing) in a unique Y-loop gene expression program.

## Discussion

The existence of the Y chromosome lampbrush-like loops of *Drosophila* has been known for the last five decades (Hess and Meyer, 1963; Meyer et al., 1961), however little is known as to how Y-loop formation and expression is regulated and whether these SC-specific structures are important for spermatogenesis. Here, we identified a Y-loop gene-specific expression program that functions in parallel to the general meiotic transcriptional program to aid in the expression and processing of the gigantic Y-loop genes. Our results suggest that genes with intron gigantism, such as the Y-loop genes and potentially other large genes such as Dystrophin, require specialized mechanisms for proper expression.

The phenotypes of *blanks* and *heph*, the two genes identified to be involved in this novel expression program, highlight two distinct steps of the Y-loop gene specific expression program (Figure 6G). Blanks was originally identified as an siRNA binding protein, but no defects in small RNA mediated silencing were observed in the testes of *blanks* mutants (Gerbasi et al., 2011; Sanders and Smith, 2011). We found that *blanks* is required for transcription of Y-loop B/*kl-3*, as nuclear transcript levels were visibly reduced in *blanks* mutants, leading to the lack of both *kl-3* mRNA granules in the cytoplasm and Kl-3 protein. As Blanks’ ability to bind RNA was previously found to be required for male fertility (Sanders and Smith, 2011), we speculate that Blanks may bind to newly synthesized nascent *kl-3* RNA, which contain megabases of satellite DNA transcripts, such that transcripts’ secondary/tertiary structures do not interfere with transcription (Zhang and Landick, 2016). It is possible that RNA polymerases, which have inherently low efficiency (Darzacq et al., 2007), might require Blanks to increase their processivity, allowing them to transcribe through repetitive DNA sequences (Fitz et al., 2018).

Heph has been implicated in a number of steps in RNA processing and translational regulation (Kafasla et al., 2012; Sawicka et al., 2008; Valcarcel and Gebauer, 1997; Wagner and Garcia-Blanco, 2001), but Heph’s exact role in the testis remained unclear despite its requirement for male fertility (Robida et al., 2010; Robida and Singh, 2003). We found that *heph* mutants fail to generate *kl-5* cytoplasmic mRNA granules even though nuclear transcript levels appeared minimally affected. This suggests that *heph* may be required for processing the long repetitive transcripts. For example, *heph* might be required to ensure proper splicing of the Y-loop gene pre-mRNAs, which is predicted to be challenging as the splicing of adjacent exons becomes exponentially more difficult as intron length increases (Shepard et al., 2009). Y-loop genes may utilize proteins like Heph to combat this challenge.

These results highlight the presence of a unique program tailored toward expressing genes with intron gigantism. Although the functional relevance of intron gigantism remains obscure, our results may provide hints as to the possible functions of intron gigantism. Even if intron gigantism did not arise to serve a specific function, once it emerges, the unique gene expression program that can handle intron gigantism must evolve to tolerate the burden of gigantic introns, as indicated by our study on *blanks* and *heph* mutants. Ultimately, the presence of a unique gene expression program for genes with gigantic introns would provide a unique opportunity to regulate gene expression. Once such systems evolve, other or new genes may start utilizing such a gene expression program to add an additional layer of complexity to the regulation of gene expression. For example, in the case of Y-loop genes, the extended time period required for the transcription of the gigantic Y-loop genes (~80-90 hours) might function as a ‘developmental timer’ for SC differentiation. Similar to this idea, it was shown that the expression of two homologous genes, *knirps* (kni) and *knirps-like* (*knrl*), is regulated by intron size during embryogenesis in *Drosophila*. Although *knrl* can perform the same function as *kni* in embryos, mRNA of *knrl* is not produced due to the presence of a relatively large (14.9kb) intron (as opposed to the small (<1kb) introns of kni), which prevents completion of *knrl* transcription during the short cell cycles of early development (Rothe et al., 1992). Thus, intron size can play a critical role in the regulation of gene expression. Alternatively, satellite DNA-containing gigantic introns could act in a manner similar to enhancers, recruiting transcriptional machinery to the Y-loop genes to facilitate expression (Shaul, 2017).

In summary, our study provides the first glimpse at how the expression of genes with intron gigantism requires a unique gene expression program, which acts on both transcription and post-transcriptional processing.

## Acknowledgements

We thank Dean Smith, Dennis McKearin, Minx Fuller, the Bloomington Stock Center and the Developmental Studies Hybridoma Bank for reagents. We thank the Yamashita laboratory and Drs. Sue Hammoud and Lei Lei for discussions and comments on the manuscript, Dr. Ruth Lehmann for helpful suggestions and Dr. Jiandie Lin for sharing equipment. This work was supported by the Howard Hughes Medical Institute (YMY) and the NIH Cellular and Molecular Biology Training Grant T32-GM007315 (JMF).

## Materials and Methods

### Fly Husbandry

All fly stocks were raised on standard Bloomington medium at 25°C, and young flies (1- to 3-day-old adults) were used for all experiments. Flies used for wild-type experiments were the standard lab wild-type strain *yw* (*y^1^w^1^*). The following fly stocks were used: *heph^2^* (BDSC:635), Df(3R)BSC687 (BDSC: 26539), *blanks^KG00084^* (BDSC:13914), Df(3L)BSC371 (BDSC:24395), p(PTT-GC)heph*^CC00664^* (BDSC:51540), *UAS-kl-3^TRiP.HMC03546^* (BDSC:53317), *UAS-blanks^TRiP.HMS00078^* (BDSC:33667), *UAS-kl-5^TRiP.HMC03747^* (BDSC:55609), and C(1)RM/C(1;Y)6, *y^1^w^1^f^1^*/0 (BDSC:9460) were obtained from the Bloomington Stock Center (BDSC). *GFP-blanks* (GFP-tagged *blanks* expressed by endogenous promoter) was a gift of Dean Smith (Sanders and Smith, 2011). *bam-gal4* was a gift of Dennis McKearin (Chen and McKearin, 2003). The *aly^2^* and *aly^5P^* stocks were a gift of Minx Fuller (Lin et al., 1996).

The Y chromosome in the *heph* deficiency strain Df(3R)BSC687 appeared to have accumulated mutations that resulted in abnormal Y-loop morphology. This Y chromosome was replaced with the *yw* Y chromosome for all experiments described in this study.

The *kl-3-FLAG* strain was constructed by Fungene (fgbiotech.com) using CRISPR mediated knock-in of a 3X-FLAG tag at the C-terminus of *kl-3* using homology-directed repair.

### RNA Fluorescent *in situ* Hybridization

All solutions used for RNA FISH were RNase free. Testes from 2–3 day old flies were dissected in 1X PBS and fixed in 4% formaldehyde in 1X PBS for 30 minutes. Then testes were washed briefly in PBS and permeabilized in 70% ethanol overnight at 4°C. Testes were briefly rinsed with wash buffer (2X saline-sodium citrate (SSC), 10% formamide) and then hybridized overnight at 37°C in hybridization buffer (2X SSC, 10% dextran sulfate (sigma, D8906), 1mg/mL E. coli tRNA (sigma, R8759), 2mM Vanadyl Ribonucleoside complex (NEB S142), 0.5% BSA (Ambion, AM2618), 10% formamide). Following hybridization, samples were washed three times in wash buffer for 20 minutes each at 37°C and mounted in VECTASHIELD with DAPI (Vector Labs). Images were acquired using an upright Leica TCS SP8 confocal microscope with a 63X oil immersion objective lens (NA = 1.4) and processed using Adobe Photoshop and ImageJ software.

Fluorescently labeled probes were added to the hybridization buffer to a final concentration of 50nM (for satellite DNA transcript targeted probes) or 100nM (for exon targeted probes). Probes against the satellite DNA transcripts were from Integrated DNA Technologies. Probes against *kl-3, kl-5, fzo*, and *Dic61B* exons were designed using the Stellaris® RNA FISH Probe Designer (Biosearch Technologies, Inc.) available online at www.biosearchtech.com/stellarisdesigner. Each set of custom Stellaris® RNA FISH probes was labeled with Quasar 670, Quasar 570 or Fluorescein-C3 (Supplemental file 1).

For strains expressing GFP (e.g. GFP-Blanks, Heph-GFP), the overnight permeabilization in 70% ethanol was omitted.

### RT-qPCR

Total RNA from testes (50 pairs/sample) was extracted using TRIzol (Invitrogen) according to the manufacturer’s instructions. 1μg of total RNA was reverse transcribed using SuperScript III® Reverse Transcriptase (Invitrogen) followed by qPCR using *Power* SYBR Green reagent (Applied Biosystems). Primers for qPCR were designed to amplify only mRNA. For average introns, one primer of the pair was designed to span the two adjacent exons.

Primers spanning large introns could only produce a PCR product if the intron has been spliced out. Relative expression levels were normalized to GAPDH and control siblings. All reactions were done in technical triplicates with at least two biological replicates. Graphical representation was inclusive of all replicates and p-values were calculated using a t-test performed on untransformed average ddct values. Primers used are listed in Supplementary file 3.

### Western blot

Testes (40 pairs/sample) were dissected in Schneider’s media at room temperature within 30 minutes, the media was removed and the samples were frozen at −80°C until use. After thawing, testes were then lysed in 200uL of 2X Laemmli Sample Buffer + βME (BioRad, 161–0737). Samples were separated on a NuPAGE Tris-Acetate gel (3–8%, 1.5mm, Invitrogen) and transferred onto polyvinylidene fluoride (PVDF) membrane (Immobilon-P, Millipore) using NuPAGE transfer buffer (Invitrogen) without added methanol. Membranes were blocked in 1X TBST (0.1% Tween-20) containing 5% nonfat milk, followed by incubation with primary antibodies diluted in 1X TBST containing 5% nonfat milk. Membranes were washed with 1X TBST, followed by incubation with secondary antibodies diluted in 1X TBST containing 5% nonfat milk. After washing with 1X TBST, detection was performed using the Pierce® ECL Western Blotting Substrate enhanced chemiluminescence system (Thermo Scientific). Primary antibodies used were anti–α-tubulin (1:2,000; mouse, monoclonal, clone DM1a; Sigma-Aldrich) and anti-FLAG (1:2,500; mouse, monoclonal, M2, Sigma-Aldrich). The secondary antibody was horseradish peroxidase (HRP) conjugated anti-mouse IgG (1:10,000; Jackson ImmunoResearch Laboratories).

### Screen for the identification of proteins involved in Y-loop gene expression

Initially, ~2200 candidate genes were selected based on gene ontology (GO) terms (e.g.. “mRNA binding”, “regulation of translation”, “spermatid development”). These genes were cross-referenced against publicly available RNAseq data sets (i.e.: FlyAtlas, modENCODE) and only those genes predicted to be expressed in the testis were selected. Additionally, candidate genes were eliminated if they are known to be involved in ubiquitous processes (e.g. general transcription factors, ribosomal subunits) or processes that are seemingly unrelated to those associated with the Y-loop genes (e.g. mitochondrial proteins, GSC/SG differentiation, mitotic spindle assembly). Finally, candidates were limited to those with available reagents for localization and/or phenotypic analysis, leaving a final list of 67 candidate genes (Supplementary file 2). If available, we first analyzed protein localization for each candidate. If candidate proteins did not localize to SCs or the Y-loops, they were not further examined. If the candidate was found to be expressed in SCs or if no localization reagents were available, then RNAi mediated knockdown or mutants were used to examine Y-loop gene expression for any deviations from the expression pattern described in Figure 1D-H and to assess fertility. As Y-loop genes are all essential for sperm maturation (Hardy et al., 1981), any genes essential for Y-loop gene expression should also be needed for fertility. All selection criteria and a summary of phenotypes observed can be found in Supplementary file 2.

### Phalloidin Staining

Testes were dissected in 1X PBS, transferred to 4% formaldehyde in 1X PBS and fixed for 30 minutes. Testes were then washed in 1X PBST (PBS containing 0.1% Triton-X) for at least 60 minutes followed by incubation with Phalloidin-Alexa546 (ThermoFisher, a22283, 1:200) antibody in 3% bovine serum albumin (BSA) in 1X PBST at 4°C overnight. Samples were washed for 60 minutes in 1X PBST and mounted in VECTASHIELD with DAPI (Vector Labs). Images were acquired using an upright Leica TCS SP8 confocal microscope with a 63X oil immersion objective lens (NA = 1.4) and processed using Adobe Photoshop and ImageJ software.

### Phase Contrast Microscopy

Seminal vesicles were dissected in 1X PBS and transferred to slides for live observation by phase contrast on a Leica DM5000B microscope with a 40X objective (NA = 0.75) and imaged with a QImaging Retiga 2000R Fast 1394 Mono Cooled camera. Images were adjusted in Adobe Photoshop.

**Supplementary Figure 1.**
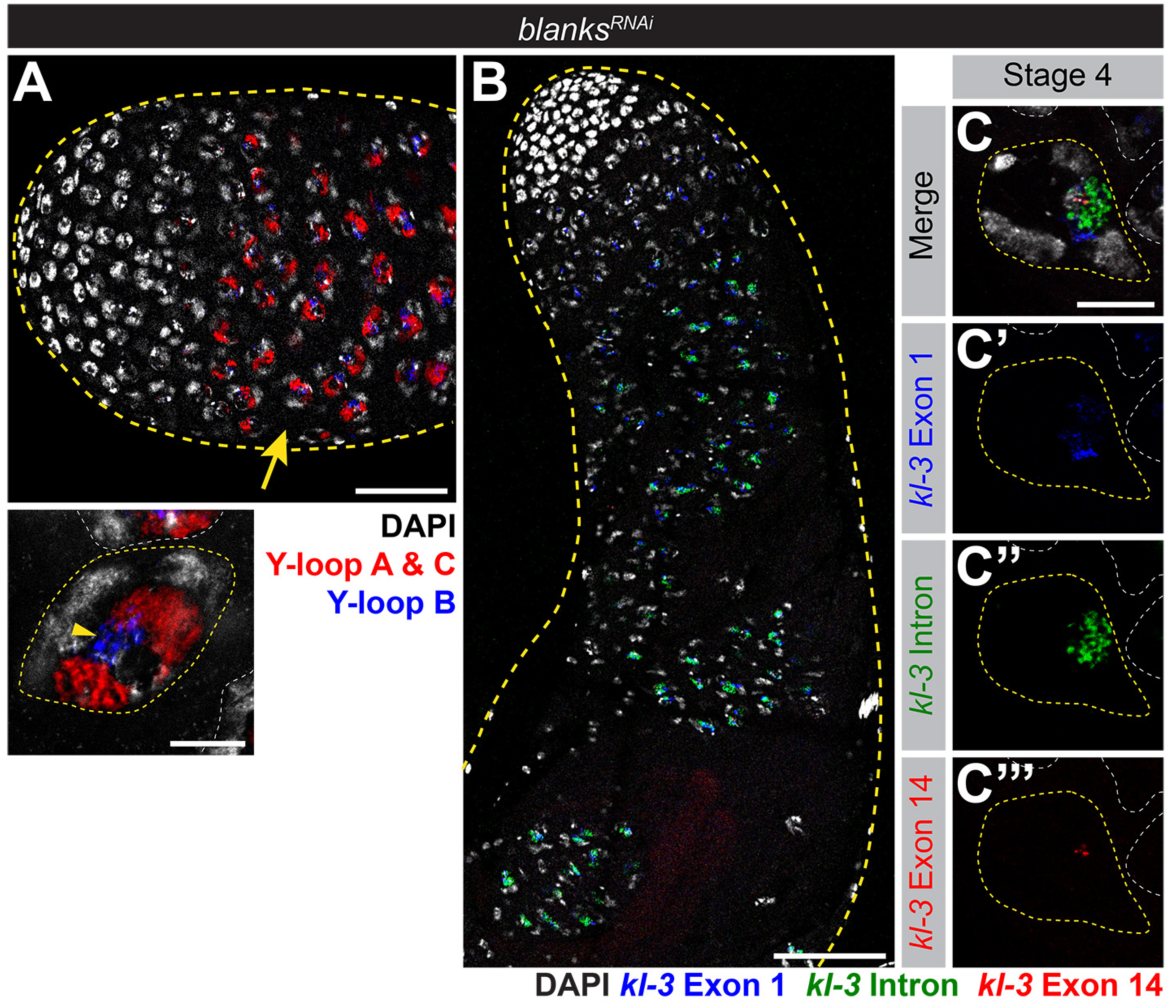
*blanks* RNAi recapitulates the phenotypes observed in *blanks* mutants. **(A)** RNA FISH against the Y-loop gene intronic transcripts in *bam-gal4>UAS-blanks^TRiP.HMS00078^* testes. Testis outline (yellow dashed line), Y-loops A and C (Cy3-(AAGAC)_6_, red), Y-loop B (Cy5-(AATAT)_6_, blue) and DAPI (white). Comparable stage SC (yellow arrow, compare to Figure 3A-B). Bar: 50 μm. High magnification image of a single SC at a comparable stage (compare to Figure 3A, B) is provided below. SC nucleus (yellow dashed line) and nuclei of neighboring cells (white dashed line). Bar: 10μm. **(B, C)** RNA FISH against *kl-3* in *bam-gal4>UAS-blanks^TRiP.HMS00078^* testes. Exon 1 (blue), *kl-3* intron (Alexa488-(AATAT)_6_, green), Exon 14 (red) and DAPI (white). **(B)** Apical third of the testis through the end of SC development (yellow dashed line). Bar: 75μm. **(C)** Single late SC nucleus (yellow dashed line). Nuclei of neighboring cells (white dashed line) and mRNA granules (yellow arrows), Bar: 10μm.

**Supplementary Figure 2.**
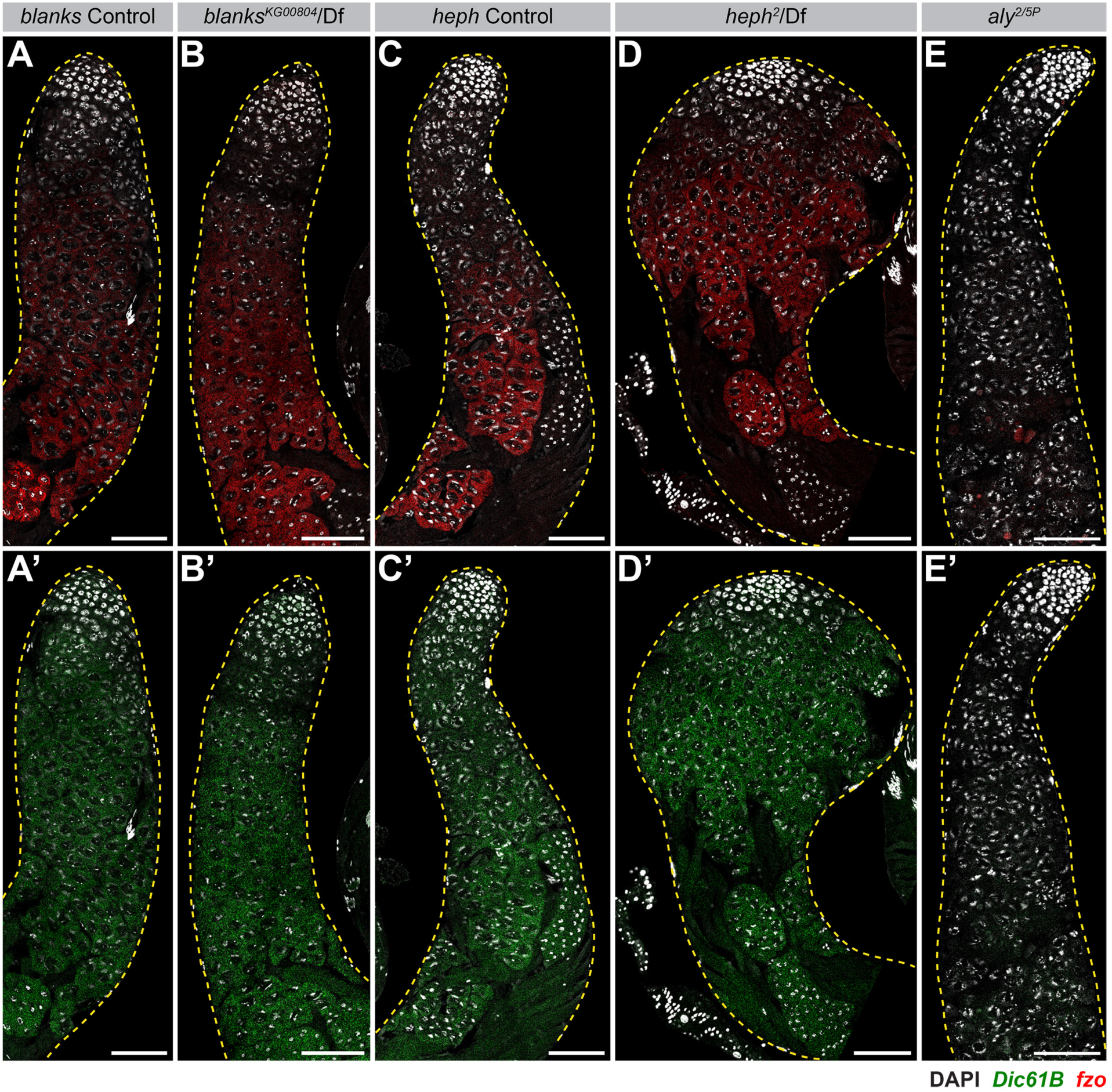
*blanks* and *heph* are not part of the meiotic transcriptional program. RNA FISH against *fzo* **(A-E)** and *Dic61B* **(A’-E’)** in *blanks* controls **(A)**, *blanks^KG00084^/Df* **(B)**, *heph* controls **(C)**, *heph****^2^***/Df **(D)**, and *aly****^2/5P^* (E)**. Apical third of the testis through the end of SC development (yellow dashed line). DAPI (white). Bar: 75μm..

## Supplementary Tables

Supplementary file 1: RNA FISH probes

Supplementary file 2: Screen

Supplementary file 3: qPCR primers

